# Peripheral Monocyte-Derived Extracellular Vesicles Establish an Immune–Brain Communication Pathway in Alzheimer’s Disease

**DOI:** 10.64898/2026.08.27.747536

**Authors:** Mingjin Wang, Weida Wang, Yanfeng Li, Zhongwu Liu, Louie Semaan, Amy Kemper, Xian Shuang Liu, Li Zhang, Michael Chopp, Zheng Gang Zhang, Yi Zhang

## Abstract

Alzheimer’s disease (AD) is increasingly recognized as a systemic disorder involving both central and peripheral immune dysfunction, yet the mechanisms by which peripheral immune cells influence neurodegeneration remain poorly understood. Here we identify a physiological extracellular vesicle (EV)-mediated route through which peripheral monocytes communicate with neurons and show that AD-associated monocyte remodeling converts this pathway into a mechanism that mediates neuronal injury. Reanalysis of single-cell transcriptomic data revealed pronounced inflammatory and EV-related transcriptional remodeling in circulating monocytes from patients with AD. Using a genetic CD63-based EV tracking mouse, we found that EVs released from peripheral Lyz2-expressing myeloid cells, including monocytes, accessed the healthy brain parenchyma and preferentially associated with neurons. EVs isolated from primary peripheral monocytes of 5xFAD mice were enriched in inflammatory cargo, including IL-1β, and markedly suppressed distal axonal growth. Neutralization of EV-associated IL-1β partially restored axonal growth, identifying IL-1β as an important mediator of EV-induced neuronal injury. Aβ stimulation reproduced key features of this pathogenic EV phenotype in RAW 264.7 macrophage-like cells and induced coordinated metabolic dysfunction and pro-inflammatory activation in primary peripheral monocytes. Moreover, repeated systemic administration of EVs from Aβ-stimulated RAW 264.7 cells accelerated behavioral and cognitive decline and reduced hippocampal synaptic integrity in 5xFAD mice without increasing cerebral amyloid plaque burden. Together, these findings reveal a peripheral monocyte-EV-neuron communication axis that operates under homeostatic conditions and can be redirected toward pathogenic signaling in AD. Targeting this EV-mediated pathway may provide a therapeutic strategy complementary to current Aβ-directed approaches.

## Introduction

Alzheimer’s disease (AD) is the most common form of dementia and is characterized by the accumulation of extracellular amyloid-β (Aβ) plaques and intracellular neurofibrillary tangles composed of hyperphosphorylated tau, accompanied by synaptic dysfunction, neuronal loss, blood–brain barrier (BBB) disruption, and progressive cognitive decline (1–3). Although reducing cerebral Aβ burden remains a major therapeutic strategy, the modest clinical benefits and treatment-associated adverse effects of anti-Aβ therapies indicate that additional mechanisms contribute to disease progression (4–6). AD is therefore increasingly viewed not solely as a brain-confined disorder, but as a systemic disease involving interactions between central pathology and peripheral immune dysfunction. Beyond its direct effects within the brain, Aβ is increasingly recognized as a potent immunomodulatory stimulus capable of altering both central and peripheral immune responses (7–9). Among peripheral immune populations, circulating monocytes are of particular interest because they contribute to Aβ uptake and clearance and undergo marked transcriptional remodeling during AD (10–12). However, peripheral monocyte infiltration into the AD brain parenchyma appears limited and is most evident at advanced disease stages (13, 14), suggesting that direct cellular entry alone is unlikely to fully explain how these cells influence neuronal function and earlier stages of disease progression.

Recent studies suggest that communication between peripheral immune cells and the central nervous system extends beyond direct cellular infiltration (15, 16). Extracellular vesicles (EVs) have emerged as key mediators of intercellular communication by transporting proteins, lipids, and nucleic acids that reflect the physiological state of their donor cells (17–19). This raises the possibility that peripheral immune cells may communicate with neural cells through EVs even in the absence of substantial cellular infiltration. In the AD brain, EVs have been implicated in the propagation of pathogenic proteins, including tau, while macrophage-derived migrasomes have recently been shown to contribute to cerebrovascular injury following Aβ stimulation (20). Circulating EVs from patients with AD have also been reported to access brain in experimental brain models and induce inflammatory responses, supporting a potential role for peripheral EVs in immune–brain communication. Because Aβ processing occurs within endosomal-lysosomal compartments that are closely linked to EV biogenesis (21, 22), Aβ may directly influence both EV production and cargo composition in peripheral monocytes. Moreover, small EVs (sEVs) can traverse the BBB (23, 24), positioning them as potential mediators of long-range brain–immune communication. Whether peripheral monocyte-derived EVs normally access neural cells and whether AD redirects this communication pathway toward pathogenic signaling, remains unknown.

In this study, we investigated whether peripheral monocyte-derived EVs provide a mechanism linking systemic immune alterations to neuronal dysfunction in AD. We employed a set of approaches including reanalysis of published human peripheral immune-cell transcriptomic datasets, genetic EV tracking in healthy mice, studies on primary monocyte and neuronal function, and longitudinal EV administration in AD mice. This integrated approach was designed to determine whether peripheral monocyte-associated EVs can access neural cells under homeostatic conditions and whether AD redirects this EV-mediated communication pathway towards pathogenic signaling. Our findings support peripheral monocyte-derived EVs as a previously underrecognized route of immune–brain communication that may contribute to AD progression.

## Results

### AD monocytes exhibit coordinated inflammatory and EV-related transcriptional remodeling

To systematically characterize peripheral immune alterations in AD, we reanalyzed a single-cell RNA-sequencing (scRNA-seq) dataset of peripheral blood mononuclear cells (PBMCs) from patients with AD and from age-matched healthy controls (25). Unsupervised Leiden clustering identified 20 transcriptionally distinct immune clusters based on canonical marker gene expression (**Supplementary Fig. 1a,b**), which were subsequently consolidated into eight major immune cell populations (**Fig. 1a**). Overall immune cell composition was largely preserved between patients with AD and healthy controls (**Fig. 1b; Supplementary Fig. 1c**). Although the abundance of circulating monocytes was modestly reduced in AD (**Supplementary Fig. 1d,e**), these cells exhibited the greatest number of differentially expressed genes among all immune populations relative to healthy controls (**Fig. 1c**). These findings indicate that peripheral immune alterations in AD are characterized primarily by pronounced transcriptional reprogramming of circulating monocytes rather than by substantial changes in immune cell composition.

**Figure 1.**
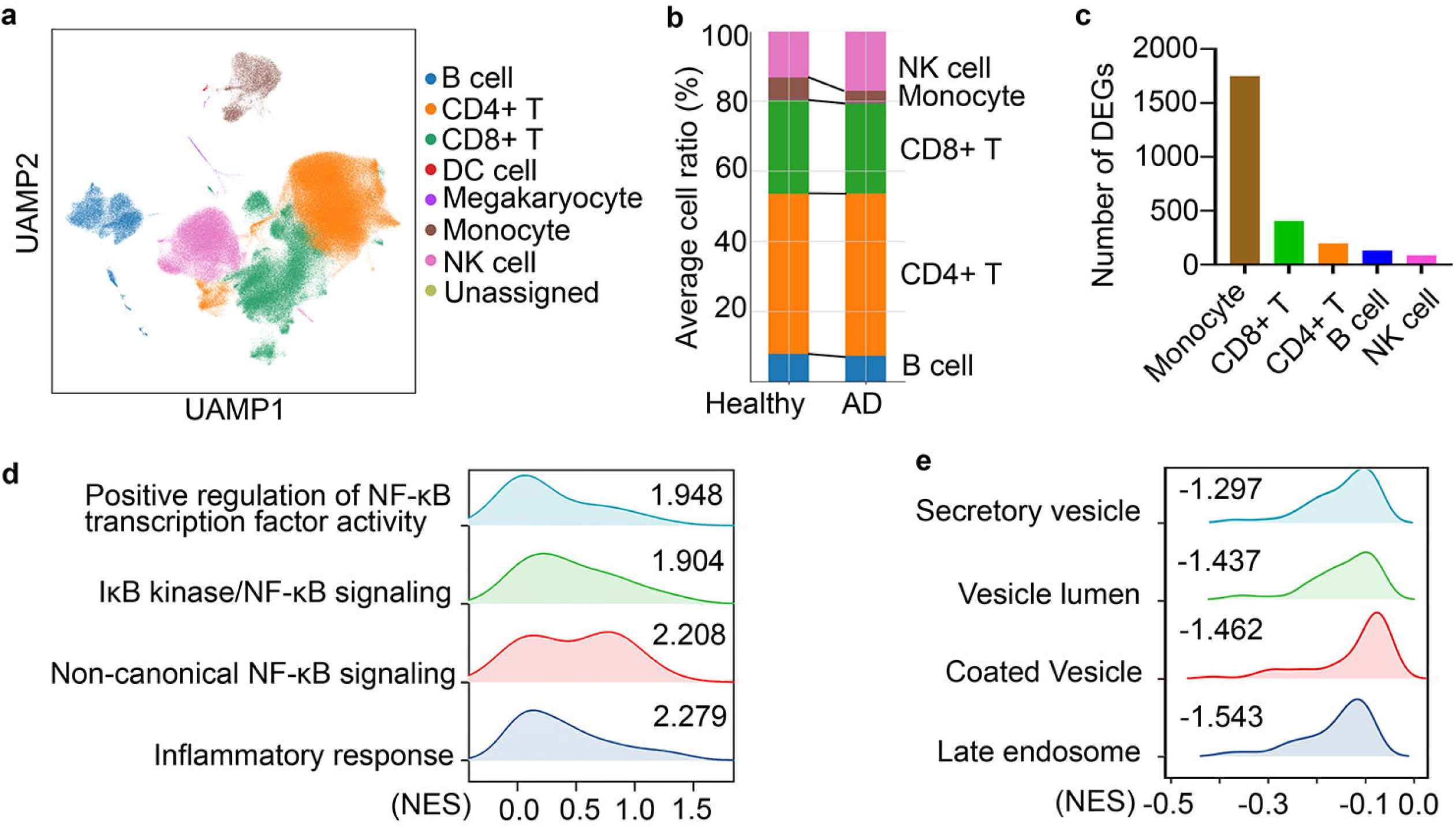
AD monocytes exhibit inflammatory and EV-biogenesis signatures. a, Uniform Manifold Approximation and Projection (UMAP) visualization of PBMCs from AD patients and age-matched healthy controls, showing eight major immune cell populations identified by canonical marker genes. b, Relative proportions of major five immune cell populations in AD and healthy control samples, showing overall preservation of peripheral immune cell composition between groups. c, Number of Differentially Expressed Genes (DEGs) identified in each major immune cell population. d, Gene Set Enrichment Analysis (GSEA) of monocytes demonstrating enrichment of inflammatory pathways. Normalized enrichment scores (NES) are listed and adjusted P <0.001. e, GSEA demonstrating negatively enrichment of extracellular vesicle-related cellular component pathways in AD monocytes. NES are listed and adjusted P <0.01. Statistical analyses are described in the Methods.

Pathway enrichment analysis further revealed activation of NF-κB-related inflammatory signaling in AD monocytes (**Fig. 1d; Supplementary Data 1**), consistent with the original report (25) and supporting a pronounced pro-inflammatory transcriptional state. Extending these findings, our focused reanalysis identified enrichment of EV- and vesicle-related pathways, including secretory vesicle, vesicle lumen, coated vesicle, and late endosome pathways (**Fig. 1e**). These pathways are associated with vesicular cargo processing, intracellular trafficking, and secretion, suggesting that EV-related cellular processes are altered in AD monocytes. Together, these findings demonstrate that circulating monocytes in AD undergo coordinated inflammatory and vesicle-associated transcriptional remodeling, providing a rationale for investigating monocyte-derived EVs as mediators of peripheral immune–brain communication.

### Peripheral monocyte–derived EVs access the healthy brain and preferentially associate with neurons

Building on the monocyte-associated transcriptional changes identified in AD, we first asked whether peripheral monocyte-associated EVs can access neural cells under homeostatic conditions. We used the Lyz2^Cre^;CD63-emGFP^l/s/l^ genetic reporter mouse(26, 27). Lyz2-Cre efficiently targets circulating monocyte populations, although its activity is not restricted exclusively to monocytes(28); in this model, Cre-mediated recombination labels CD63-positive small EVs released from Lyz2-expressing myeloid cells and enables their endogenous tracking in vivo (**Fig. 2a**).

**Figure. 2.**
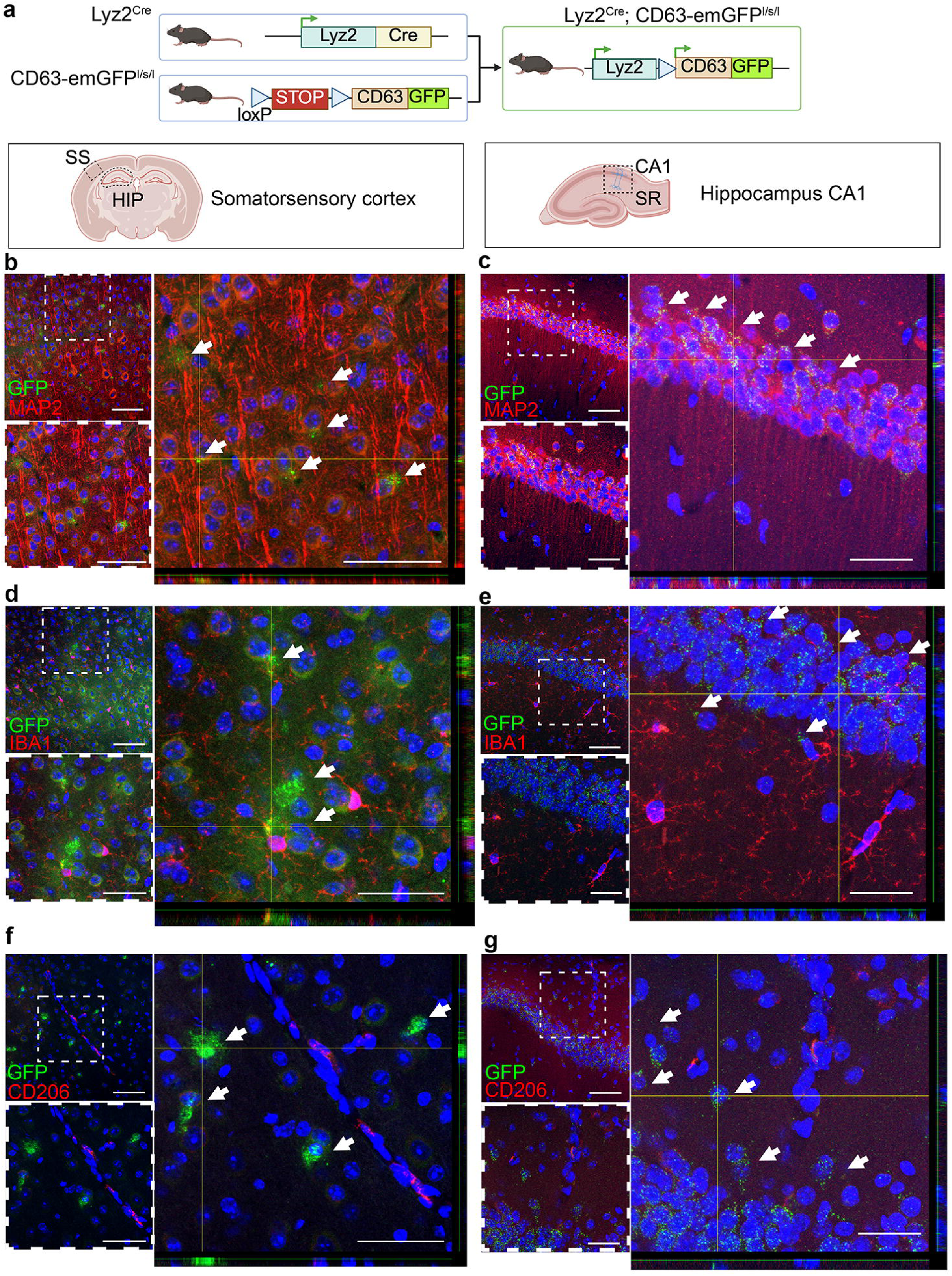
Peripheral monocyte EVs enter the brain and preferentially associate with neurons. a, Schematic of the Lyz2^Cre^;CD63-emGFP^l/s/l^ reporter mouse used for endogenous labeling of CD63-positive EVs released from Lyz2-expressing monocyte-lineage cells. Brain regions analyzed are indicated, including the somatosensory cortex (SS) and hippocampal CA1 region. b-g, Representative confocal images of GFP-positive monocyte-derived EVs (green) in the somatosensory cortex (b,d,f) and hippocampal CA1 (c,e,g), co-stained with MAP2-positive neurons (b,c), IBA1-positive microglia (d,e), or CD206-positive perivascular macrophages (f,g) (red). Dashed boxes indicate regions shown at higher magnification, with corresponding three-dimensional reconstructions in the right panels. GFP-positive EVs were frequently observed in close apposition to neuronal cell bodies and neurites, but showed limited spatial association with microglia or perivascular macrophages. Arrowheads indicate representative GFP-positive EVs. Scale bars, 50 μm (low magnification) and 25 μm (high magnification).

Confocal analysis of coronal brain sections from healthy reporter mice showed that lysozyme-positive myeloid cells were localized primarily to the choroid plexus and vascular and meningeal interfaces, with little evidence of their presence within the brain parenchyma (**Supplementary Fig. 2a**). This distribution is consistent with the preferential localization of peripheral and border-associated myeloid cells at CNS interfaces under steady-state conditions(13, 29). In parallel, monocytes isolated from the peritoneal cavity exhibited robust GFP expression, and GFP-positive EVs were readily detected in EV preparations derived from these cells (**Supplementary Fig. 2b,c**), confirming effective labeling of peripheral monocytes and their released EVs in this reporter system.

Despite the restricted localization of lysozyme-positive cells, abundant GFP-positive EV puncta were detected throughout the somatosensory cortex (**Fig. 2b,d,f**) and hippocampal CA1 region (**Fig. 2c,e,g**), indicating that EVs released from Lyz2-expressing peripheral myeloid cells can access and distribute within the brain parenchyma. Three-dimensional reconstruction of confocal z-stacks showed that GFP-positive EV puncta were frequently located in close apposition to MAP2-positive neuronal cell bodies and neurites (**Fig. 2b,c**). In contrast, comparatively little spatial association was observed with IBA1-positive microglia (**Fig. 2d,e**) or CD206-positive perivascular macrophages (**Fig. 2f,g**), suggesting preferential association with neurons. No GFP signal was detected in brain sections from wild-type littermates (**Supplementary Fig. 3**).

Together, these findings show that EVs released from peripheral Lyz2-expressing myeloid cells, including monocytes, access the healthy brain parenchyma and preferentially associate with neurons without extensive infiltration of their parental cells. These observations identify a potential EV-mediated route of peripheral immune–brain communication under homeostatic conditions. We therefore next asked whether AD alters the molecular composition and neuronal effects of peripheral monocyte-derived EVs.

### AD reprograms peripheral monocytes to release inflammatory EVs that suppress neuronal growth

Having established that peripheral monocyte-associated EVs can access the healthy brain and associate with neurons, we next asked whether AD redirects this EV-mediated communication pathway toward pathogenic signaling. We therefore examined the molecular composition and neuronal effects of EVs released from primary peripheral monocytes isolated from 6-month-old 5xFAD mice (AD-Ms) and age-matched wild-type littermates (WT-Ms), followed by isolation and characterization of their corresponding EVs (AD-M-EVs and WT-M-EVs) (**Fig. 3a,b**, **Supplementary Fig 4**). EV concentrations are comparable between groups: 2.43 × 10^10^ ± 6.73×10^8^particles/ml for AD-M-EVs and 2.87 × 10^10^ ± 1.04×10^9^ particles/ml for WT-M-EVs.

**Fig. 3.**
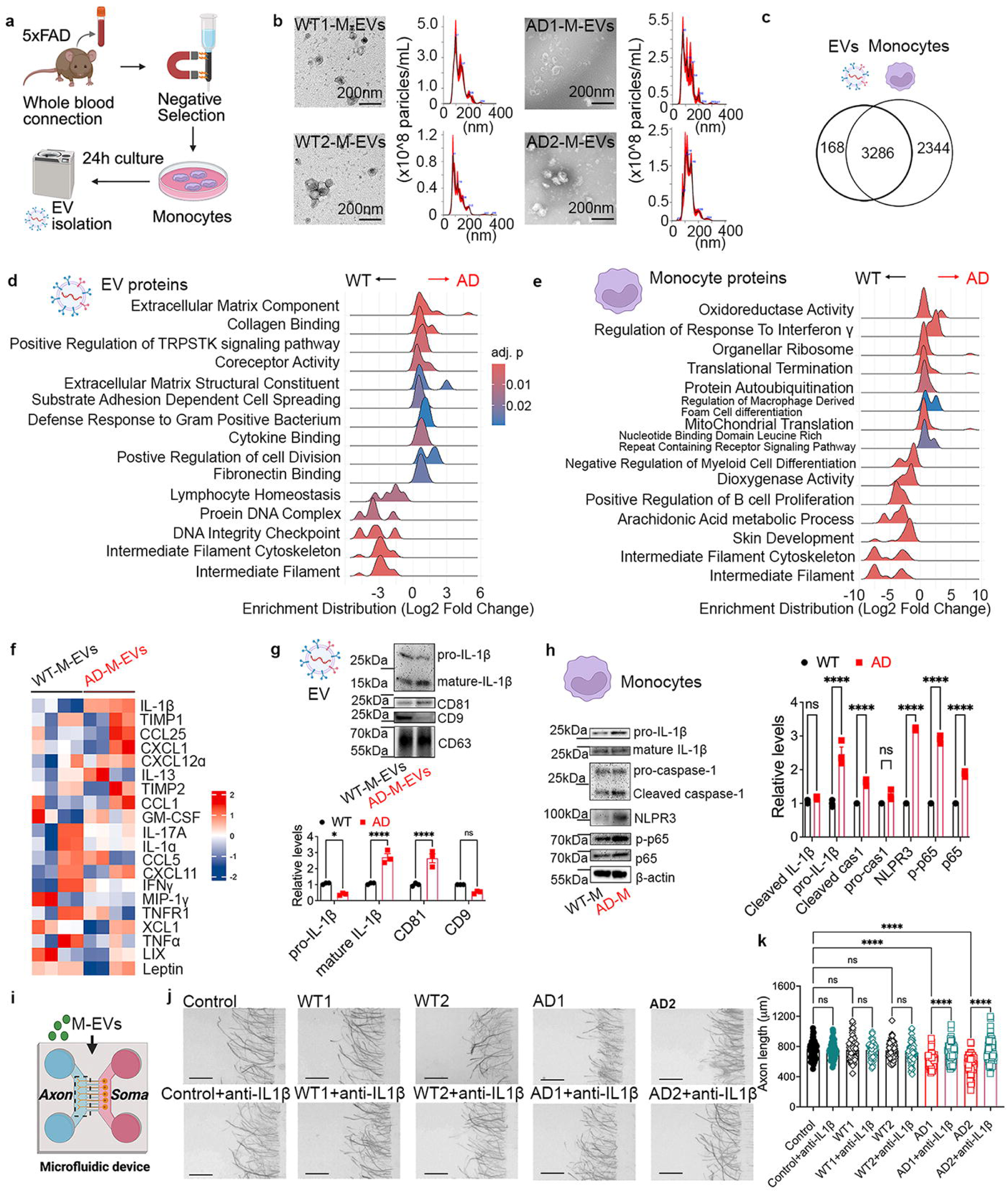
AD monocytes produce inflammatory EVs that suppress neuronal growth. a, Schematic of the experimental workflow. Ly6C^+^ peripheral monocytes were isolated from whole blood of 6-month-old wild-type (WT) and 5xFAD (AD) mice by negative selection, cultured, and EVs were collected for downstream analyses. b, Representative transmission electron microscopy images and nanoparticle tracking analysis of EVs isolated from WT and AD monocytes. EVs exhibited typical cup-shaped morphology and comparable size distributions. c, Venn diagram showing overlap between proteins identified by mass spectrometry in monocytes and their corresponding EVs. d-e, Gene ontology enrichment analysis of differentially expressed proteins in EVs (d) and parental monocytes (e) comparing AD and WT samples. f, Cytokine array analysis demonstrating altered inflammatory cytokine profiles in EVs isolated from WT and AD monocytes. g-h, Representative Immunoblot validation and corresponding quantification data show EV-associated (g) and parental monocyte inflammatory proteins (h). i, Schematic of the microfluidic neuron culture platform used to evaluate the effects of monocyte-derived EVs on axonal growth. EVs were applied to both the somatic and axonal compartment. j, Representative images of distal axons following treatment with EVs isolated from WT or AD monocytes in the presence or absence of IL-1β neutralization. k, Quantification of axonal length following EV treatment. Data are presented as mean ± s.e.m. Scale bars in i, 200 μm. Statistical analyses are described in the Methods.

To determine how AD alters monocyte-derived EVs, we performed unbiased proteomic profiling of both parental monocytes and their corresponding EVs. Mass spectrometry identified a substantial overlap between monocyte and EV proteomes while revealing distinct EV-enriched protein subsets (**Fig. 3c, Supplementary Data 2**). Compared with WT-M-EVs, AD-M-EVs exhibited significant enrichment of proteins associated with extracellular matrix organization, proteoglycan-containing extracellular matrix, extracellular structure organization, and cytokine binding (**Fig. 3d**), indicating that AD selectively remodels EV cargo toward inflammatory and tissue-remodeling functions rather than globally altering protein release. Proteomic analysis of the parental monocytes also revealed broad AD-associated cellular remodeling, including pathways related to responses to interferon-γ, mitochondrial and organellar translation, NLR-containing receptor signaling, arachidonic acid metabolism, and regulation of myeloid-cell differentiation (**Fig. 3e**). Together, these findings indicate that the altered EV phenotype arises in the context of coordinated inflammatory and metabolic remodeling of the EV-producing monocytes.

To validate these findings, cytokine array analysis demonstrated broad elevation of pro-inflammatory cytokines and chemokines in AD-M-EVs compared with WT-M-EVs (**Fig. 3f, Supplementary Data 3**). Among these, IL-1β showed one of the most prominent increases. Consistent with this observation, immunoblot analysis showed that mature IL-1β was significantly increased in both AD monocytes and their corresponding EVs (**Fig. 3g,h**). In parental AD monocytes, pro-IL-1β was also elevated, together with increased cleaved caspase-1, NLRP3, and phosphorylated p65, consistent with enhanced IL-1β production, inflammasome-associated processing, and NF-κB activation. In contrast, pro-IL-1β was not increased in AD-M-EVs, suggesting preferential enrichment of the mature IL-1β form within the EV compartment. AD-M-EVs also displayed increased CD81 and reduced CD9 expression relative to WT-M-EVs (**Fig. 3g**), suggesting that AD not only alters EV cargo but also changes the molecular composition of monocyte-derived EV populations.

Using a compartmentalized microfluidic neuron culture system (30, 31) permitting neurons to grow their axons into distal axonal chamber, we next examined whether AD-M-EVs directly affect axonal growth (**Fig. 3i**). Treatment with AD-M-EVs markedly reduced distal axonal outgrowth compared with WT-M-EVs (**Fig. 3j,k**). Preincubation of EVs with an IL-1β-neutralizing antibody (Abcam, Cat. No. ab2105; 1:200; RRID:AB_302842) significantly attenuated this inhibitory effect and partially restored axonal growth, indicating that EV-associated IL-1β is an important mediator of AD-M-EV-induced axonal injury.

Together, these findings demonstrate that AD reprograms peripheral monocyte-EVs to be enriched with inflammatory proteins and these pathogenic cargo directly suppress axonal growth.

### A**β** induces neurotoxic EV production in RAW 264.7 macrophage-like cells

Having identified an inflammatory and neurotoxic EV phenotype in primary peripheral monocytes from 5xFAD mice, we next asked whether Aβ alone was sufficient to reproduce this phenotype. To isolate the direct effects of Aβ from the complex systemic environment of AD and establish a standardized EV-production model for subsequent mechanistic and in vivo studies, we used RAW 264.7 murine macrophage-like cells. Cells were exposed to 100 nM oligomeric Aβ_₁₋₄₂_, a concentration previously used to examine cellular responses to soluble Aβ oligomers in vitro ^30,31^. EVs released from control and Aβ-treated RAW 264.7 cells were subsequently isolated and characterized (**Supplementary Fig. 5**).

Transmission electron microscopy (TEM) and nanoparticle tracking analysis (NTA) demonstrated that EVs isolated from Aβ-treated RAW 264.7 cells (Aβ-R-EVs) showed morphology, size distribution, and expression of canonical EV markers comparable to those of control EVs (c-R-EVs), while displaying a modest increase in particle concentration (control: 5.17 × 10^10^± 8.61 × 10^8^ particles/mL versus Aβ: 6.21 × 10^10^ ± 2.14 × 10^9^ particles/mL; **Supplementary Fig. 5a-d**). Single-particle phenotyping by Exoview (32, 33) further showed comparable expression of canonical EV markers and myeloid-associated surface markers, suggesting that Aβ modestly increased EV release without substantially altering the overall surface-marker profile of the EV population (**Supplementary Fig. 5e**). Consistent with our observations in primary monocytes isolated from 5xFAD mice, immunoblot analysis showed that Aβ increased both pro-IL-1β and mature IL-1β in RAW 264.7 cells, whereas their corresponding EVs exhibited increased mature IL-1β without a significant change in pro-IL-1β (**Fig. 4a**). This pattern closely recapitulated the IL-1β processing profile observed in primary AD monocytes and their EVs, indicating that Aβ alone is sufficient to reproduce a key feature of the inflammatory EV phenotype associated with AD.

**Fig 4.**
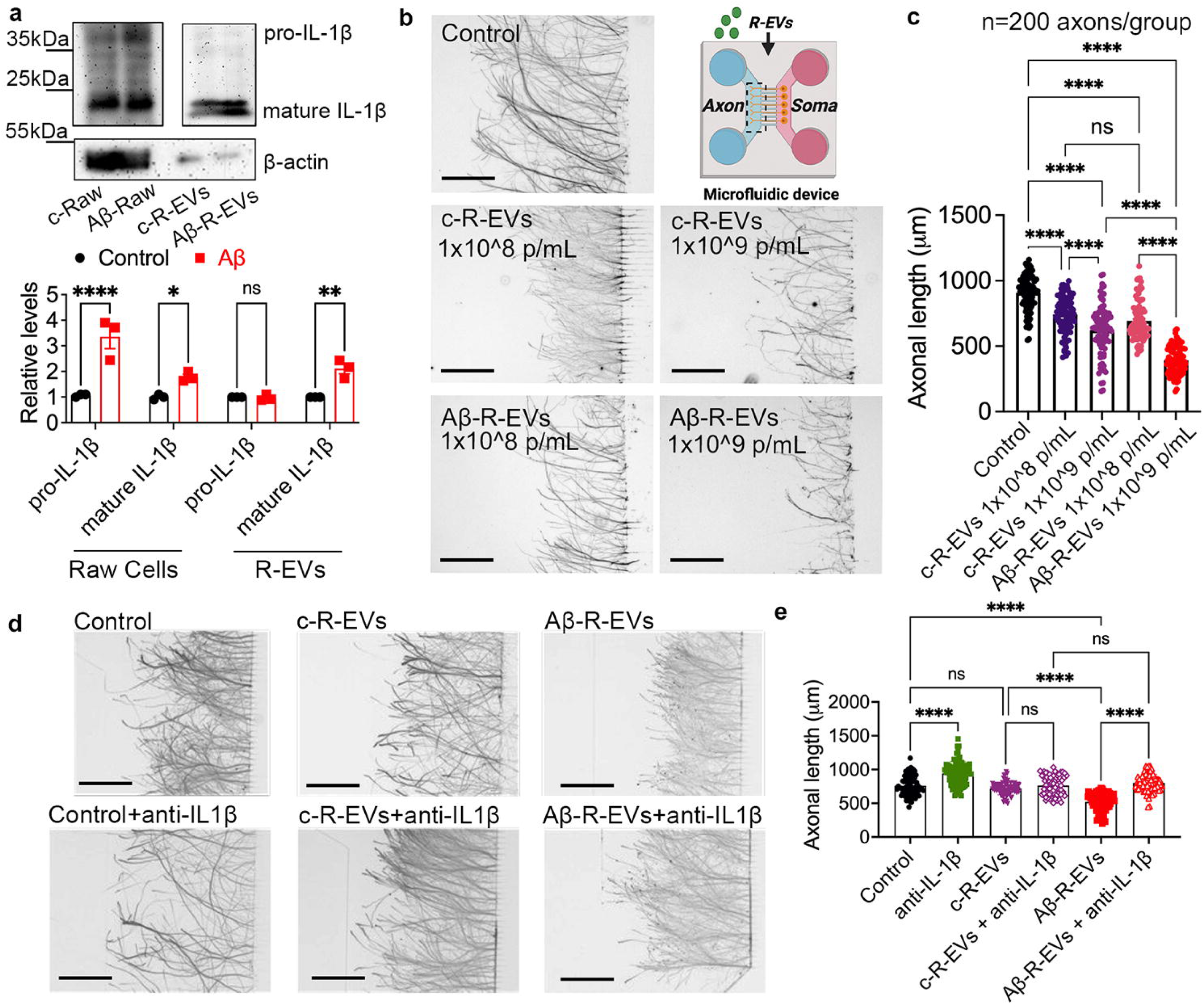
Amyloid-β reprograms monocytes to produce neurotoxic EVs. a, Representative immunoblots and quantification showing IL-1β expression in RAW264.7 monocytes (cells) and corresponding EVs following stimulation with oligomeric Aβ_1-42_. b-c, Representative axonal images (b) and quantification data (c) showing axonal length of cortical neurons treated with control monocyte-derived EVs (c-R-EVs) or EVs isolated from Aβ-stimulated monocytes (Aβ-R-EVs) at the indicated particle concentrations. Upper right, schematic of the microfluidic device. d-e, Representative axonal images (d) and quantification data (e) showing axonal length of cortical neurons following treatment with c-R-EVs or Aβ-R-EVs in the presence or absence of an IL-1β-neutralizing antibody. Data are presented as mean ± s.e.m. Scale bars, b,d, 200 μm. Statistical analyses are described in the Methods.

Using the same compartmentalized microfluidic device approach, we next examined the effects of Aβ-R-EVs on axonal growth. Aβ-R-EVs markedly suppressed distal axonal outgrowth compared with control EVs (c-R-EVs) (**Fig. 4b**), and this effect increased with EV concentration (**Fig. 4b,c**). In contrast, direct exposure of neuronal axons to 100 nM oligomeric Aβ_1-42_ did not significantly affect axonal extension under the same experimental conditions (**Supplementary Fig. 5f,g**). These findings indicate that the observed axonal injury is mediated by EVs released from Aβ-stimulated myeloid cells rather than by direct exposure to soluble Aβ.

To determine whether EV-associated IL-1β contributed to this neurotoxic effect, Aβ-R-EVs were preincubated with the IL-1β-neutralizing antibody before neuronal treatment. IL-1β neutralization significantly, but incompletely restored axonal growth (**Fig. 4d,e**), identifying EV-associated IL-1β as an important functional mediator while suggesting that additional EV cargo components also contribute to the neurotoxic phenotype.

Collectively, these findings demonstrate that Aβ alone is sufficient to induce RAW 264.7 macrophage-like cells to release inflammatory and neurotoxic EVs that suppress axonal growth, thereby reproducing key features of the pathogenic EV phenotype identified in AD-associated primary monocytes.

### A**β** induces metabolic dysfunction and pro-inflammatory activation in primary peripheral monocytes

Having established in RAW 264.7 cells that Aβ stimulation is sufficient to generate neurotoxic EVs, we next determined whether Aβ induces a comparable pathogenic state in primary peripheral monocytes. Proteomic analysis of monocytes from 5xFAD mice had revealed enrichment of inflammatory and metabolic pathways, including interferon-γ responses, mitochondrial translation, NLR-containing receptor signaling, and arachidonic acid metabolism (**Fig. 3e**). We therefore asked whether Aβ alone was sufficient to reproduce key features of this AD-associated monocyte phenotype. Because cellular metabolism is closely linked to innate immune activation and EV production, primary peripheral monocytes isolated from healthy C57/BL6 mouse whole blood were exposed to Aβ_₁₋₄₂_ and analyzed for metabolic and inflammatory responses..

Seahorse metabolic flux analysis revealed marked impairment of both oxidative phosphorylation and glycolysis in Aβ-treated monocytes (**Fig. 5a–d**). Aβ significantly reduced oxygen consumption rate and extracellular acidification rate, indicating broad suppression of mitochondrial respiration and glycolytic activity, and identifying Aβ as a potent metabolic stressor of peripheral monocytes.

**Fig 5.**
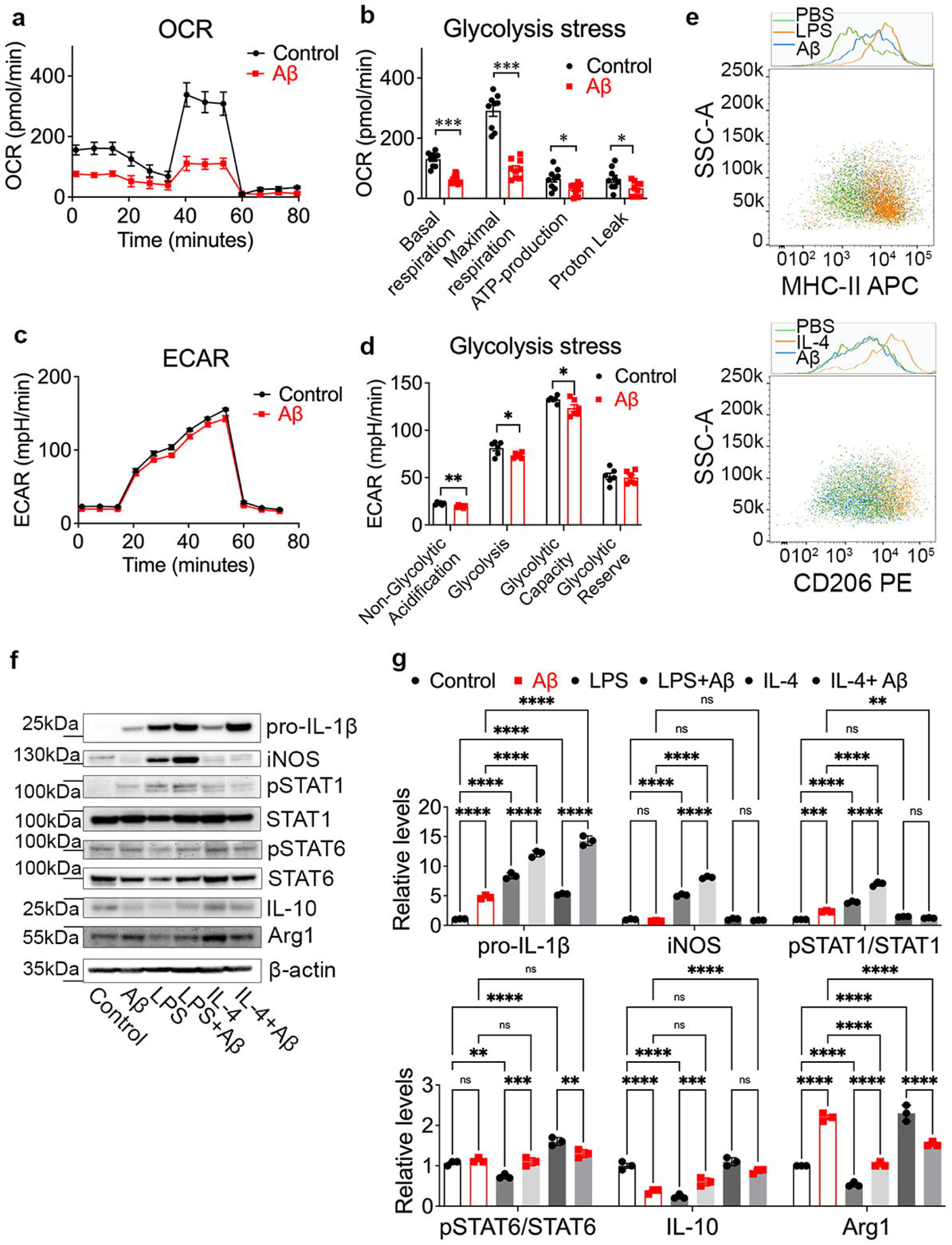
Aβ induces metabolic dysfunction and pro-inflammatory polarization in monocytes. a, Seahorse extracellular flux analysis showing oxygen consumption rate (OCR) in Ly6c+ peripheral monocytes following treatment with vehicle control or oligomeric Aβ1-42. b, Quantification of mitochondrial respiratory parameters derived from OCR measurements, including basal respiration, maximal respiration, ATP production, and proton leak. c, Seahorse extracellular acidification rate (ECAR) measurements showing glycolytic activity in control and Aβ-treated monocytes. d, Quantification of glycolytic parameters, including non-glycolytic acidification, glycolysis, glycolytic capacity, and glycolytic reserve. e, Flow cytometric analysis of monocyte polarization following Aβ stimulation. Representative histograms and contour plots showing expression of the M1-associated marker MHC-II and the M2-associated marker CD206. Lipopolysaccharide (LPS) and IL-4 were included as positive controls for M1 and M2 polarization, respectively. f-g, Representative immunoblot images (f) and quantification data (g) showing inflammatory polarization markers in monocytes treated with vehicle, Aβ, LPS, LPS+Aβ, IL-4, or IL-4+Aβ. Data are presented as mean ± s.e.m. Statistical analyses are described in the Methods.

Because metabolic remodeling is closely coupled to innate immune activation, we next examined whether Aβ alters monocyte inflammatory polarization. Flow cytometry showed increased surface expression of the pro-inflammatory marker MHC-II following Aβ stimulation, whereas CD206 expression remained unchanged (**Fig. 5e**). Immunoblotting further revealed increased IL-1β, inducible nitric oxide synthase, and phosphorylated STAT1, accompanied by reduced IL-10, Arg1, and phosphorylated STAT6 (**Fig. 5f,g**), supporting a shift toward a pro-inflammatory phenotype. Compared with the classical pro-inflammatory stimulus LPS, Aβ induced a similar inflammatory program without further enhancing the LPS response, while partially suppressing IL-4-induced anti-inflammatory signaling. These findings suggest that Aβ promotes a sustained pro-inflammatory state in primary peripheral monocytes.

Together, these results demonstrate that Aβ induces coordinated metabolic dysfunction and pro-inflammatory activation in primary peripheral monocytes, recapitulating key features of the inflammatory and metabolic remodeling identified in monocytes from 5xFAD mice and providing a physiologically relevant cellular context for the inflammatory EV phenotype observed in the RAW 264.7 myeloid model..

### A**β**-stimulated myeloid cell-derived EVs accelerate cognitive decline and synaptic dysfunction in vivo

Having established that Aβ stimulation generates neurotoxic EVs in a controlled myeloid model and induces a comparable pathogenic state in primary peripheral monocytes, we next determined whether Aβ-R-EVs were sufficient to accelerate AD-associated functional decline in vivo. RAW 264.7-derived EVs were used to provide a standardized and reproducible EV preparation for repeated systemic administration.

Beginning at 3 months of age, female 5xFAD mice were administered Aβ-R-EVs intraperitoneally every other day until 6 months of age. Mice receiving PBS or control RAW 264.7-derived EVs (c-R-EVs) served as controls. Behavioral function was assessed longitudinally at 3, 4, and 6 months of age, and neuropathological analyses were performed at 6 months (**Fig. 6a**).

**Fig 6.**
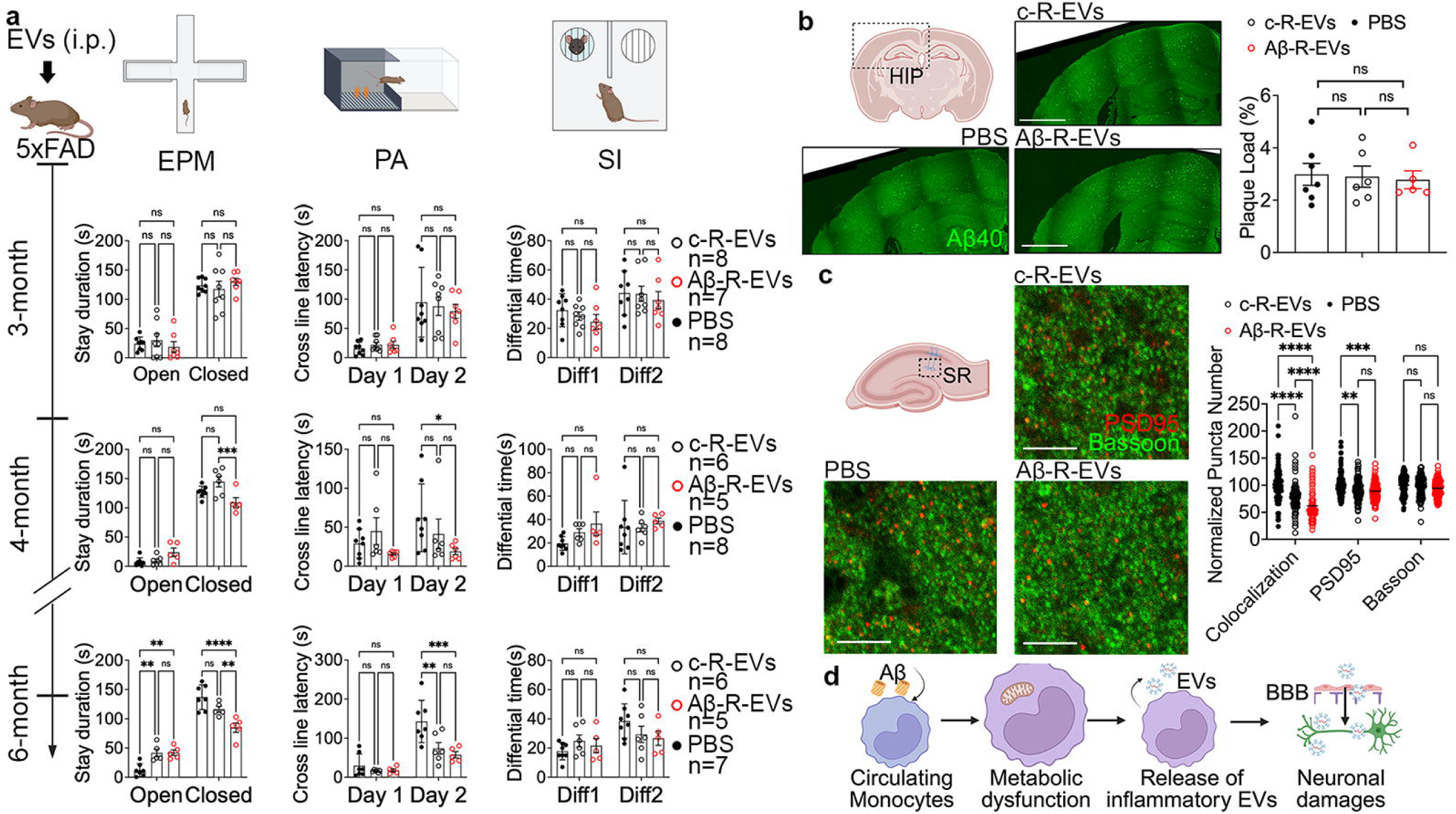
Aβ-induced monocyte-derived EVs accelerate Alzheimer’s disease progression in vivo. a, Experimental design and longitudinal behavioral assessment. Female 5xFAD mice beginning at 3 months of age received repeated intravenous administration of PBS, control c-R-EVs, or Aβ-R-EVs. Behavioral testing was performed longitudinally at 3, 4, and 6 months using the elevated plus maze (EPM), passive avoidance (PA), and social interaction (SI) assays. b, Representative immunofluorescence images and quantification of Aβ plaque burden in the somatosensory cortex (SS) and hippocampus (HIP). c, Representative images and quantification of hippocampal synaptic puncta labeled by PSD95 and Bassoon. d, Proposed model illustrating peripheral monocyte-mediated brain–immune communication in Alzheimer’s disease. Aβ exposure induces metabolic and inflammatory reprogramming of circulating monocytes, promoting the release of inflammatory extracellular vesicles enriched in pathogenic cargo. These EVs cross the blood–brain barrier, preferentially associate with neurons, and contribute to synaptic dysfunction and cognitive decline. Data are presented as mean ± s.e.m. Statistical analyses are described in the Methods.

Mice receiving Aβ-R-EVs developed progressively greater anxiety-like behavior than PBS- or c-R-EV-treated mice, as indicated by reduced time spent in the open arms of the elevated plus maze (**Fig. 6a**). Passive avoidance testing further revealed impaired memory retention in Aβ-R-EV-treated mice at 6 months, whereas social interaction remained unchanged (**Fig. 6a**). These findings indicate that Aβ-R-EVs selectively exacerbate anxiety-related and memory-associated behavioral deficits in 5xFAD mice.

To determine whether these functional deficits were accompanied by changes in AD neuropathology, we quantified amyloid plaque burden and synaptic integrity. In a separate tracking experiment, systemically administered fluorescently labeled EVs were detected in the brain parenchyma (**Supplementary Fig. 6**). Repeated administration of Aβ-R-EVs did not significantly alter amyloid plaque burden in either the somatosensory cortex or hippocampus compared with PBS or c-R-EV treatment (**Fig. 6b**). In contrast, Aβ-R-EV-treated mice exhibited marked reductions in PSD95-positive postsynaptic puncta and Bassoon/PSD95-colocalized puncta within the hippocampal CA1 stratum radiatum (**Fig. 6c**), a region containing Schaffer collateral synapses critical for CA3-to-CA1 transmission and hippocampus-dependent memory (34, 35). These structural changes were consistent with the progressive behavioral deficits observed in Aβ-R-EV-treated mice, suggesting that Aβ-R-EVs accelerate functional decline by compromising hippocampal synaptic integrity rather than by increasing amyloid plaque accumulation.

Together, these findings demonstrate that EVs released from Aβ-stimulated myeloid cells are sufficient to accelerate behavioral and cognitive decline and synaptic dysfunction in 5xFAD mice without exacerbating amyloid plaque burden, supporting the concept that a peripheral EV-mediated communication route can be redirected toward pathogenic signaling in AD.

## Discussion

AD has traditionally been viewed as a brain-centered disorder driven by amyloid-β (Aβ) accumulation, tau pathology, and neuroinflammation (1–3). However, accumulating evidence suggests that peripheral immune dysregulation also contributes to disease progression, although the mechanisms linking peripheral immunity to neuronal dysfunction remain poorly understood (36, 37). In the present study, we identify an EV-mediated route (**Fig. 6d**) through which peripheral monocytes may communicate with neurons and show that this pathway can be redirected toward pathogenic signaling in AD. Our results show that AD-associated monocytes release inflammatory EVs with neurotoxic activity. Aβ reproduced key features of this phenotype by inducing inflammatory remodeling of peripheral monocytes and the release of IL-1β-enriched EVs. These EVs suppressed axonal growth and accelerated behavioral and synaptic dysfunction in 5xFAD mice. Together, our findings support peripheral monocyte-derived EVs as a long-range communication mechanism through which systemic immune alterations can amplify neuronal injury in AD.

One of the major conceptual advances of this study is the identification of a peripheral EV pathway that is anatomically present under homeostatic conditions but can acquire pathogenic properties in AD. Although circulating monocytes have long been implicated in AD, their direct contribution remains controversial because their infiltration into the brain parenchyma appears limited and is most evident at advanced disease stages(14, 38). Recent evidence that age-dependent replacement of hippocampal microglia by monocyte-like cells occurs in the human brain further highlights the increasing contribution of peripheral myeloid cells to brain homeostasis and disease with aging(39). Our findings suggest that monocyte-derived EVs provide an alternative mechanism by which peripheral immune alterations can affect neurons without requiring extensive cellular entry into the brain. Using a genetic EV reporter mouse, we demonstrate that monocyte-derived EVs are present throughout the brain parenchyma and preferentially associate with neurons rather than resident microglia or perivascular macrophages. These observations establish the anatomical feasibility of peripheral monocyte-to-neuron EV communication but do not by themselves define its physiological function. Our subsequent findings suggest that AD-associated changes in monocytes can redirect this pre-existing route toward inflammatory and neurotoxic signaling. These results extend recent reports demonstrating that macrophage-derived migrasomes contribute to cerebrovascular injury and that neuronal EVs propagate pathogenic proteins such as tau(20, 40). Collectively, these studies support the emerging concept that EVs are active mediators of immune-brain communication during neurodegeneration.

Our study further identifies Aβ as an extracellular factor capable of redirecting this peripheral EV communication pathway. While Aβ has primarily been investigated for its direct neurotoxic effects within the brain(41), our findings demonstrate that it also profoundly reprograms peripheral monocytes. Exposure to Aβ induced coordinated metabolic and inflammatory remodeling characterized by impaired oxidative phosphorylation, reduced glycolytic capacity, mitochondrial dysfunction, and activation of pro-inflammatory signaling pathways (42, 43). These changes were accompanied by increased IL-1β expression and the release of inflammatory EVs. However, our data do not establish that metabolic dysfunction directly causes EV cargo remodeling, and intervention studies targeting specific metabolic pathways will be required to define this relationship. Moreover, Aβ is unlikely to be the only circulating factor capable of altering monocyte function; cytokines, damage-associated molecules, lipids, and other AD-associated signals may cooperate with or act independently of Aβ. Thus, our findings establish that Aβ alone is sufficient to induce key features of the pathogenic EV phenotype, rather than identifying it as the sole upstream driver.

The pathogenic activity of monocyte-derived EVs appears to depend, at least in part, on their inflammatory cargo. Proteomic and cytokine analyses revealed selective enrichment of extracellular matrix-associated proteins and inflammatory mediators in EVs derived from AD monocytes, indicating that AD-associated conditions remodel EV composition rather than simply altering EV release. Among these factors, IL-1β emerged as a prominent mediator. Neutralization of EV-associated IL-1β significantly restored axonal growth, supporting a causal role for IL-1β in EV-induced neuronal injury. Because the neutralizing antibody was applied to intact EV preparations, this effect suggests that a biologically active pool of IL-1β is exposed on, or otherwise accessible at, the EV surface. The incomplete rescue further indicates that additional surface-associated or luminal EV cargo contributes to the neurotoxic phenotype. Notably, AD monocytes showed enhanced IL-1β processing and inflammasome activation, whereas their EVs showed increased mature IL-1β without a corresponding increase in pro-IL-1β, suggesting preferential association of the mature cytokine with EVs. A similar pattern was reproduced following Aβ stimulation of RAW 264.7 cells, further supporting Aβ as a sufficient trigger for this inflammatory EV phenotype. Given the established role of IL-1 signaling in chronic neuroinflammation and synaptic dysfunction(44). these findings identify surface-accessible EV-associated IL-1β as an important link between peripheral monocyte activation and neuronal injury. Further studies using protease-protection, membrane-permeabilization, and EV-cargo fractionation approaches will be required to define the relative contributions of surface-associated and intraluminal factors.

Importantly, our findings support the in vivo relevance of this mechanism. Repeated systemic administration of EVs released from Aβ-stimulated RAW 264.7 cells accelerated anxiety-related and memory-associated behavioral deficits and reduced hippocampal synaptic integrity in 5xFAD mice. These effects occurred without a significant increase in amyloid plaque burden, indicating that inflammatory EVs can worsen neuronal and synaptic dysfunction without further enhancing cerebral amyloid deposition. Consistent with this concept, recently identified mitochondrial plaques arising from defective mitophagy and lysosomal dysfunction represent an additional Aβ-associated but partly independent pathological process that may persist despite amyloid clearance(6). Thus, these EVs are not merely biomarkers of peripheral immune activation but can function as pathogenic mediators under the experimental conditions tested. This dissociation between functional decline and plaque burden may help explain why reducing cerebral Aβ alone produces incomplete clinical benefit. Anti-Aβ therapies target amyloid accumulation but may not fully interrupt downstream peripheral immune responses or EV-mediated inflammatory signaling. Targeting EV production, pathogenic cargo loading, or EV interactions with recipient neural cells may therefore provide a complementary therapeutic strategy, although the specificity and safety of such approaches will require careful evaluation because EV-mediated communication also serves normal physiological functions.

Our findings also have implications for emerging immune cell–based therapies for neurodegenerative diseases. Recent studies have shown that transplantation of hematopoietic progenitors or functional myeloid cells can restore microglial activity and ameliorate AD-related pathology in experimental models(45, 46), yet efficient engraftment of peripheral cells into the adult brain remains limited by the BBB and the self-renewing nature of resident microglia. Our results raise the possibility that therapeutic benefits of peripheral immune-cell replacement may not require extensive cellular engraftment. Instead, monocyte-derived EVs, which readily access the brain parenchyma, may function as key mediators of peripheral immune-to-brain communication. Although the EVs examined here were pathogenic, the existence of a peripheral EV route to neural cells also raises the possibility that appropriately engineered monocytes or EVs could deliver reparative signals to the brain. This possibility remains speculative and will require direct testing of therapeutic cargo, biodistribution, cell specificity, and long-term safety.

Several limitations should be acknowledged. First, the Lyz2^Cre^;CD63-emGFP^l/s/l^ reporter labels a broad Lyz2-expressing myeloid compartment and is not restricted exclusively to circulating monocytes. Although we confirmed reporter activity in isolated peripheral monocytes and their EVs, contributions from other Lyz2-lineage cells cannot be excluded. In addition, the reporter experiments were performed in healthy mice and establish endogenous EV access and cellular association under homeostatic conditions rather than AD-specific trafficking. Second, detection of EV signals within the brain parenchyma does not resolve the molecular route of BBB passage, and close apposition to neuronal structures does not necessarily demonstrate internalization or intracellular cargo delivery. Third, RAW 264.7 cells are murine macrophage-like cells rather than primary circulating monocytes. They provided a standardized platform for EV production, mechanistic manipulation, and repeated in vivo administration, but they may not fully reproduce the biology or EV composition of primary monocytes. Finally, although Aβ induced both metabolic dysfunction and inflammatory EV production, a direct causal relationship between these processes remains to be established. Validation using EVs isolated directly from patients with AD and monocyte-specific genetic models will be important for establishing the translational and lineage-specific relevance of this pathway.

## Methods

### Human single-cell RNA sequencing analysis

Publicly available scRNA-seq data from PBMCs of patients with AD and age-matched healthy controls were obtained from the Gene Expression Omnibus (GEO; accession GSE226602) as previously reported(25). Raw count matrices were analyzed using the Scanpy single-cell analysis pipeline implemented in Python(47). Cells expressing fewer than 200 detected genes, greater than 5,000 genes, or more than 10% mitochondrial transcripts were excluded, and genes detected in fewer than three cells were removed. After library-size normalization to 10,000 counts per cell and log-transformation, cells were clustered using the Leiden algorithm and visualized by UMAP. Immune cell identities were assigned based on the expression of established canonical marker genes, including LYZ, S100A8, S100A9 (monocytes), CD3D, IL7R (CD4⁺ T cells), CD3D, NKG7 (CD8⁺ T cells), MS4A1, CD79A (B cells), NKG7, GNLY (NK cells), FCER1A, CLEC10A (dendritic cells), and PPBP (megakaryocytes). Differential gene expression between AD and control monocytes was performed using the Wilcoxon rank-sum test implemented in Scanpy, and significantly altered genes were subjected to GSEA.

GSEA was performed using the pre-ranked algorithm in GSEApy (v.4.1.0)(48), with genes ranked by the log₂ fold change multiplied by −log₁₀(Benjamini–Hochberg-adjusted P value) from the Wilcoxon rank-sum test (Scanpy). General pathway enrichment was assessed against the MSigDB C5 ontology collection (v.2023.2.Hs)(49). Pathways with FDR q < 0.25 were considered significantly enriched.

### Animals

Female C57BL/6J mice (Jax# 000664) and breeding pairs of 5xFAD mice (Jax# 034848), Lyz2^Cre^ mice (Jax# 004781), CD63-emGFP^l/s/l^ (Jax# 036865) were obtained from the Jackson Laboratory (Bar Harbor, Maine, USA) and maintained under environment approved by Henry Ford Institutional Animal Care and Use Committee (IACUC). Lyz2^Cre^ mice were crossed with CD63-emGFP^l/s/l^ to generate Lyz2^Cre^;CD63-emGFP^l/s/l^ mice for endogenous labeling of monocyte-derived EVs by using the strategies we previously published(27). All animal procedures were approved by the IACUC of Henry Ford Hospital (Protocol No. 1482 and 1511) and were performed in accordance with NIH guidelines.

Unless otherwise indicated, female mice aged 6 months were used. For longitudinal EV administration experiments, female 5xFAD mice were enrolled at 3 months of age following genotyping. 5xFAD mice exhibiting obvious developmental abnormalities, severe illness, or non-AD-related neurological deficits before study initiation were excluded according to the guideline of The Model Organism Development and Evaluation for Late-onset Alzheimer’s Disease (MODEL-AD) Consortium (50). All remaining mice were randomly assigned to experimental groups and analyzed at 6 months of age.

### Preparation of oligomeric A**β**_1-42_

Oligomeric Aβ_1-42_was prepared according to a published protocol with minor modifications (51). Briefly, synthetic human Aβ_1-42_ peptide (MilliporeSigma, Cat. No. PP69) was dissolved in hexafluoroisopropanol (HFIP) to dissociate pre-existing aggregates and incubated for 2 h at room temperature. HFIP was evaporated under vacuum to generate a peptide film, which was stored at −80 °C until use. Immediately before experiments, the peptide film was resuspended in anhydrous dimethyl sulfoxide (DMSO) to 5 mM, sonicated for 10 min, diluted in sterile PBS, and incubated at 4 °C for 12 h to generate soluble oligomers. Aliquots were stored at −80 °C and thawed only once before use..

### Cell culture

Primary peripheral monocytes were isolated from whole blood collected from adult mice using the EasySep Mouse Monocyte Isolation Kit (STEMCELL, Cat. No.19861) according to the manufacturer’s instructions. Monocyte purity (> 90%) was confirmed by flow cytometry and immunoblotting. Purified monocytes were cultured in RPMI-1640 medium supplemented with 10% fetal bovine serum (FBS; MilliporeSigma, Cat. No. 12306C), 1% penicillin-streptomycin, and maintained at 37 °C in a humidified incubator with 5% CO₂. Where indicated, monocytes were stimulated with 100 nM oligomeric Aβ_1-42_ for the indicated times before EV isolation, Seahorse metabolic flux analysis, flow cytometry, or immunoblotting.

RAW 264.7 murine macrophage cells (ATCC TIB-71, RRID: CVCL_0493) were cultured in Dulbecco’s modified Eagle’s medium (DMEM; Thermo Fisher Scientific, Cat. No. 11966092) supplemented with 10% FBS, 1% penicillin-streptomycin. Cells were stimulated with 100 nM oligomeric Aβ_1-42_ for the indicated times before EV isolation.

### EV isolation

Cells were cultured in growth medium containing EV-depleted FBS (System Biosciences, Cat. No. EXO-FBS-250A-1) for 24 h before conditioned medium collection. Small EVs were then isolated by differential ultracentrifugation as previously described (52, 53). Briefly, conditioned medium was centrifuged sequentially at 300 × g for 10 min, 2,000 × g for 20 min, and 10,000 × g for 30 min to remove cells and debris. Small EVs were collected by ultracentrifugation at 100,000 × g for 2 h at 4°C using a Beckman Coulter Optima XPN-100 with a SW32Ti rotor. EV pellets were resuspended in sterile PBS for downstream analyses. Particle concentration, size distribution and morphology were determined immediately using NTA and TEM, as previously described (52, 53).

### Nanoparticle tracking analysis (NTA)

EV size distribution and particle concentration were measured using a NanoSight NS300 (Malvern Panalytical) equipped with a 488 nm laser as previously described (52, 53). Three videos of 60s each were recorded for every sample and analyzed using NTA software version 3.3 Dev Build 3.3.104.

### Transmission electron microscopy (TEM)

To examine EV morphology, EV suspensions were adsorbed onto carbon-coated copper grids (Electron Microscopy Sciences, Cat. No. FCF400-Cu), negatively stained with 2% uranyl acetate, and imaged using a JEOL JEM-1400 Flash transmission electron microscope (JEOL, Tokyo, Japan) operating at 80 kV, as previously described (52–54).

### ExoView analysis

To further characterize EV surface marker expression, single-particle phenotyping was performed using the ExoView R100 platform (Unchained Labs, Pleasanton, CA, USA) according to the manufacturer’s instructions and our previous studies(32, 33). EVs were captured on antibody-coated chips from the Exosome Mouse Tetraspanin Kit (Unchained Labs, Cat. No. 251-1046), containing capture antibodies against CD81, CD9, together with rat IgG and hamster IgG isotype controls. Captured EVs were subsequently immunolabeled with AF647-conjugated anti-CD11b (BioLegend, Cat. No. 101220, RRID:AB_493546) and AF488-conjugated anti-CD36 (BioLegend, Cat. No. 336231, RRID:AB_2814229). Images were acquired and analyzed using ExoView Analyzer (Unchained Labs).

### Proteomic analysis

Proteomic analysis was performed as previously described with minor modifications (55, 56). Briefly, EVs or their corresponding parental monocytes were lysed in RIPA buffer containing protease inhibitors. Equal amounts of protein (30 μg) were reduced, alkylated, and digested with sequencing-grade trypsin using the S-Trap sample preparation protocol. Peptides were separated by reversed-phase liquid chromatography and analyzed by tandem mass spectrometry on a Q Exactive mass spectrometer (Thermo Fisher Scientific) at the Wayne State University Proteomics Core Facility. Raw data were processed using Proteome Discoverer v2.4 (Thermo Fisher Scientific) with the Sequest search engine against the UniProt mouse protein database. Protein identifications were filtered at a 1% false discovery rate (FDR) and used for subsequent bioinformatic analyses.

### Cytokine array

Inflammatory cytokines in EV lysates were profiled using the Mouse Inflammation Cytokine Array C1 (RayBiotech, Cat. No. AAM-INF-1-8) according to the manufacturer’s instructions. Briefly, equal amounts of EV protein (20 μg) were incubated with cytokine array membranes overnight at 4 °C. Following incubation with detection antibodies, chemiluminescent signals were visualized and imaged. Spot intensities were quantified using ImageJ and normalized to the positive control spots on each membrane.

### Western blotting

Cells and EVs were lysed in RIPA buffer supplemented with protease and phosphatase inhibitor cocktails. Equal amounts of protein (2–10 μg) were separated by SDS–PAGE on 4–12% precast gels (Thermo Fisher Scientific, NP0322BOX) and transferred onto PVDF membranes. Membranes were blocked with 0.2% iBlock solution (Thermo Fisher Scientific), T2015) for 1 h at room temperature and incubated overnight at 4 °C with the following primary antibodies: IL-1β (Abcam, ab2105, 1:1000, RRID:AB_302842), Caspase-1 (Santa Cruz Biotechnology, sc-56036, 1:1000, RRID:AB_781816), NLRP3 (Cell Signaling Technology, 15101, 1:1000, RRID:AB_2722591), p65 (MilliporeSigma/Chemicon, MAB3026, 1:1000, RRID:AB_2178887), phospho-p65 (Cell Signaling Technology, 3033, 1:1000, RRID:AB_331284), STAT1 (Cell Signaling Technology, 9172, 1:1000, RRID:AB_2198300), phospho-STAT1 (Cell Signaling Technology, 9177, 1:1000, RRID:AB_2197983), STAT6 (Cell Signaling Technology, 9362, 1:1000, RRID:AB_2271211), phospho-STAT6 (Cell Signaling Technology, 9364, 1:1000, RRID:AB_2271227), iNOS (Abcam, ab15323, 1:1000, RRID:AB_1977477), Arginase-1 (Cell Signaling Technology, 93668, 1:1000, RRID:AB_2800207), IL-10 (Abcam, ab9969, 1:1000, RRID:AB_308826), CD63 (Santa Cruz Biotechnology, sc-5275, 1:1000, RRID:AB_627877), CD81 (BioLegend, 349502, 1:1000, RRID:AB_10643417), CD9 (Abcam, ab223052, 1:1000, RRID:AB_2922392), Calnexin (BioLegend, 669401, 1:1000, RRID:AB_2728519), and β-actin (Abcam, ab6276, 1:5000, RRID:AB_2223210). Membranes were subsequently incubated with HRP-conjugated secondary antibodies for 1 h at room temperature. Immunoreactive bands were visualized and imaged using enhanced chemiluminescence (Thermo Fisher Scientific, A38554). Band intensities were quantified using ImageJ and normalized to β-actin or the corresponding total protein, as appropriate. Uncropped immunoblot images are provided in **Supplementary Data 4**.

### Seahorse metabolic flux analysis

Cellular metabolic function was assessed by measuring the oxygen consumption rate (OCR) and extracellular acidification rate (ECAR) using an Agilent Seahorse XF96 Extracellular Flux Analyzer according to a published protocol with minor modifications(57). Briefly, primary peripheral monocytes (5×10^4^ cells/well) were seeded into Seahorse XF96 cell culture microplates, and sensor cartridges were hydrated and calibrated according to the manufacturer’s instructions. Prior to analysis, cells were equilibrated in Seahorse XF assay medium.

For the Mito Stress Test, OCR was measured following sequential injections of oligomycin (1 μM), FCCP (1 μM), and rotenone/antimycin A (0.5 μM each). For the Glycolysis Stress Test, ECAR was measured following sequential injections of glucose (10 mM), oligomycin (1 μM), and 2-deoxy-D-glucose (2-DG, 50 mM). Basal respiration, ATP-linked respiration, maximal respiration, glycolysis, glycolytic capacity, and glycolytic reserve were calculated using Wave software (Agilent Technologies).

### Flow cytometry

Primary peripheral monocytes were collected, washed with ice-cold FACS buffer composed by PBS containing 2% Bovine Serum Albumin (BSA), and incubated with anti-mouse CD16/32 Fc Block (BioLegend, 101301, RRID:AB_312800) for 10 min at 4 °C. Cells were subsequently stained with fluorophore-conjugated antibodies against MHC-II (BioLegend, Cat. No. 107613, RRID:AB_313328) and CD206 (BioLegend, Cat. No. 141707, RRID:AB_10896057) for 1 h at 4 °C in the dark. Following washing, cells were resuspended in FACS buffer and analyzed on a Sony MA900 Cell Sorter (Sony Biotechnology). Data were analyzed using FlowJo (v10, BD Biosciences). Monocytes were identified based on forward and side scatter characteristics, and doublets and debris were excluded prior to analysis.

### Primary neuronal culture and microfluidic assay

Primary cortical neurons were isolated from embryonic day 17–18 Sprague–Dawley rat embryos as previously described (30). Neurons were seeded into Xona microfluidic chambers (SND450, Xona Microfluidics) precoated with poly-D-lysine and laminin and maintained in Neurobasal medium supplemented with B27, GlutaMAX, and penicillin-streptomycin (Thermo Fisher Scientific). Extracellular vesicles (EVs) were added to the somatic compartment at the indicated particle concentrations. For IL-1β neutralization experiments, EVs were preincubated with an anti-IL-1β neutralizing antibody (Abcam, Cat. No. ab2105, RRID:AB_302842) for 1 h at room temperature before neuronal treatment. At the indicated time points, cultures were fixed with 4% paraformaldehyde, washed with phosphate-buffered saline (PBS), and immunostained with an antibody against βIII-tubulin (TUJ1) (BioLegend, Cat. No. 801202, RRID:AB_2313773). Images were acquired using an Olympus FV1200 confocal microscope, and axonal length and branching were quantified using ImageJ.

### Immunohistochemistry and confocal microscopy and image analysis

Brains were fixed in 4% paraformaldehyde, processed for formalin-fixed paraffin embedding, and sectioned coronally at 30 μm and followed with IHC as we previously described(52). The following primary antibodies were used: GFP (Abcam, Cat. No. ab290, RRID:AB_2313768; 1:200), MAP2 (Abcam, Cat. No. ab32454, RRID:AB_776174; 1:200), IBA1 (Abcam, Cat. No. ab5076, RRID:AB_2224402; 1:100), CD206 (R&D Systems, Cat. No. AF2535, RRID:AB_2063012; 1:100), lysozyme (Thermo Fisher Scientific, Cat. No. MA5-32154, RRID:AB_2809443; 1:100), PSD95 (Cell Signaling Technology, Cat. No. 3409, RRID:AB_1264242; 1:100), Bassoon (Abcam, Cat. No. ab82958, RRID:AB_1860018; 1:100), and Aβ_1-42_ (BioLegend, Cat. No. 805509, RRID:AB_2783381; 1:100). Confocal z-stack images were acquired using an Olympus FV1200 laser-scanning confocal microscope under identical acquisition settings across experimental groups. Image analysis and three-dimensional reconstruction were performed using Fiji/ImageJ as previously described(52, 54).

For quantification of amyloid plaque load, anatomically matched sections containing the somatosensory cortex and hippocampus were analyzed using ImageJ as previously described. Briefly, regions of interest were segmented using a fixed fluorescence-intensity threshold applied uniformly. Plaque load was calculated as the percentage of each region occupied by Aβ-positive staining. Measurements from 3 sections per animal, separated by approximately 100 μm, were averaged to generate one value for each animal.

Synaptic density and integrity in the CA1 stratum radiatum were quantified as previously described with minor modifications(58). Briefly, confocal z-stack images were analyzed using Imaris (Bitplane) and ImageJ. The numbers of Bassoon-positive, PSD95-positive, and colocalized puncta were quantified, with colocalized puncta considered representative of excitatory synapses. Measurements from 3 images obtained from 3 sections per animal were averaged for statistical analysis.

### In vivo EV administration

Three-month-old female 5xFAD mice were randomly assigned to receive PBS, control EVs (c-R-EVs), or Aβ stimulated EVs (Aβ-R-EVs). EVs were administered intraperitoneally at a dose of 1×10^10^ particles per injection every other day and 3 times per week from 3 to 6 months of age. Control mice received an equivalent volume of PBS on the same schedule.

For exogenous EV tracking, purified EVs were labeled using the RNASelect Green Fluorescent Staining Kit (Thermo Fisher Scientific, Cat. No. S32703) as previously described(53). Labeled EVs were administered in vivo 2 h before tissue collection, and EV-associated fluorescence was examined by immunofluorescence microscopy in 30-μm brain cryosections.

### Behavioral testing

Investigators performing behavioral testing and histological analyses were blinded to treatment allocation. Behavioral assessments were conducted longitudinally at 3, 4, and 6 months of age according to published protocols with minor modifications(59–62). For the elevated plus maze, mice were placed in the center of the apparatus and allowed to explore freely for 5 min. The time spent in the open and closed arms was quantified. For the passive avoidance test, mice underwent habituation and training on day 1, during which entry into the dark chamber was paired with a mild foot shock. Memory retention was assessed 24 h later by measuring the latency to enter the dark chamber. For the three-chamber social interaction test, mice were allowed to explore an apparatus containing a novel mouse confined beneath a wire enclosure in one side chamber and an empty enclosure in the opposite chamber. Sociability was assessed during a 10-min session by quantifying the time spent interacting with the novel mouse versus the empty enclosure.

### Statistical analysis

Data are presented as mean ± s.e.m. Statistical analyses were performed using GraphPad Prism v10. Comparisons between two groups were analyzed using unpaired two-tailed Student’s t-tests. Comparisons among multiple groups were performed using one-way or two-way ANOVA followed by Tukey’s multiple-comparison test. Longitudinal behavioral data were analyzed using repeated-measures two-way ANOVA where appropriate. P < 0.05 was considered statistically significant.

