## Supplementary Figures for "Peripheral Monocyte-Derived Extracellular Vesicles Establish an Immune–Brain Communication Pathway in Alzheimer’s Disease"

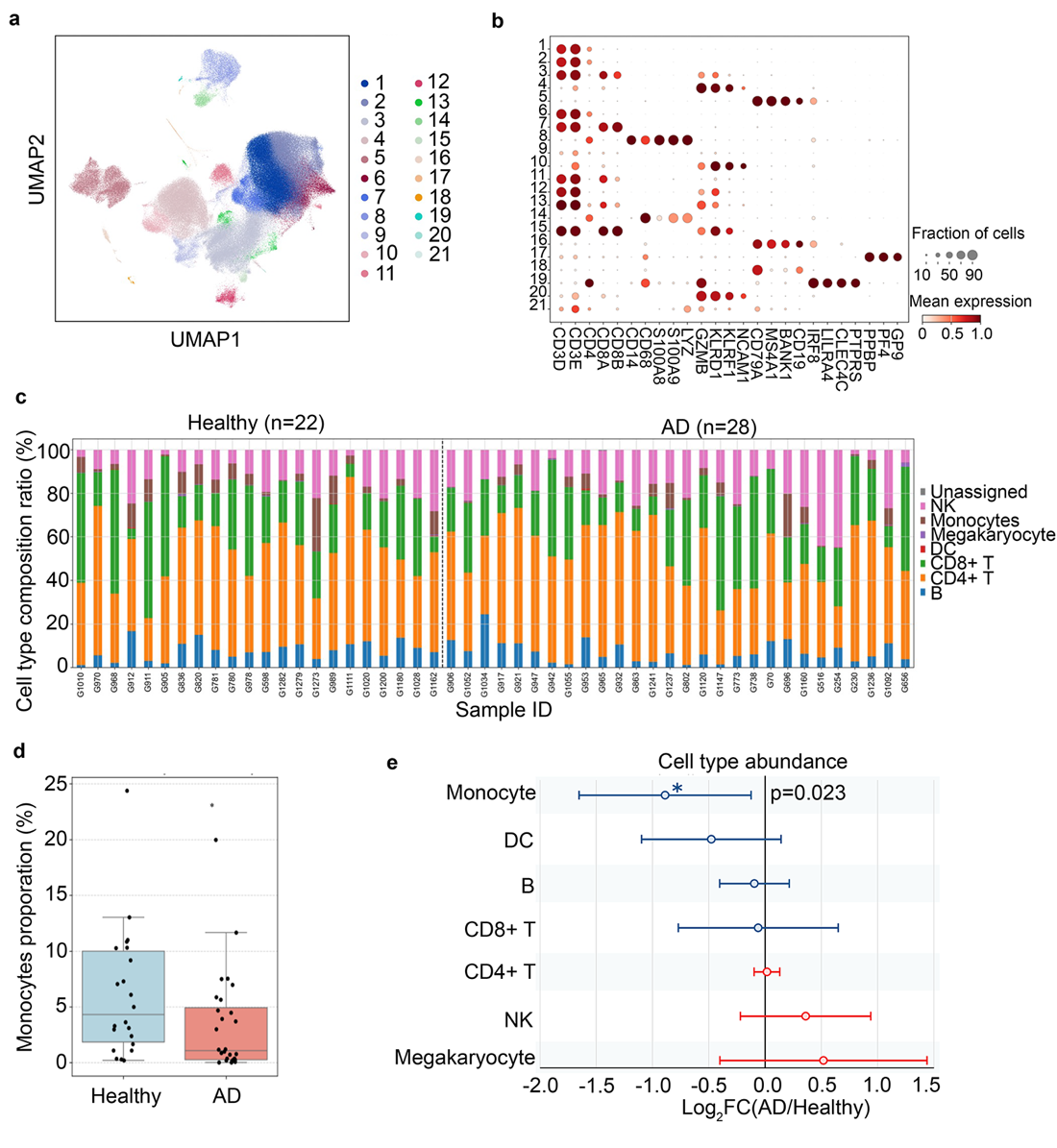

**Supplementary Figure. 1** Single-cell transcriptomic analysis identifies peripheral monocytes as the major immune population altered in AD.

a, Unsupervised Leiden clustering of PBMCs from AD patients and age-matched healthy controls identified 21 transcriptionally distinct cell clusters. Clusters were visualized by UMAP.

b, Dot plot showing expression of canonical marker genes used to annotate individual clusters into major immune cell populations. Dot size indicates the fraction of cells expressing each marker, and color intensity represents the average normalized expression level.

c, Relative abundance of major immune cell populations across individual healthy control and AD samples. Overall immune cell composition was largely preserved between groups.

d, Quantification of circulating monocyte proportions in healthy controls and AD patients. Monocyte abundance was modestly reduced in AD compared with healthy controls. Each dot represents one individual sample. Box plots indicate the median and interquartile range, with whiskers representing the minimum and maximum values.

e, Summary of differential cell-type abundance analysis comparing AD and healthy control samples. Monocytes were the only major immune population exhibiting a significant change in abundance, whereas other immune cell populations were not significantly altered.

Data are presented as mean ± s.e.m. Statistical analyses are described in the Methods

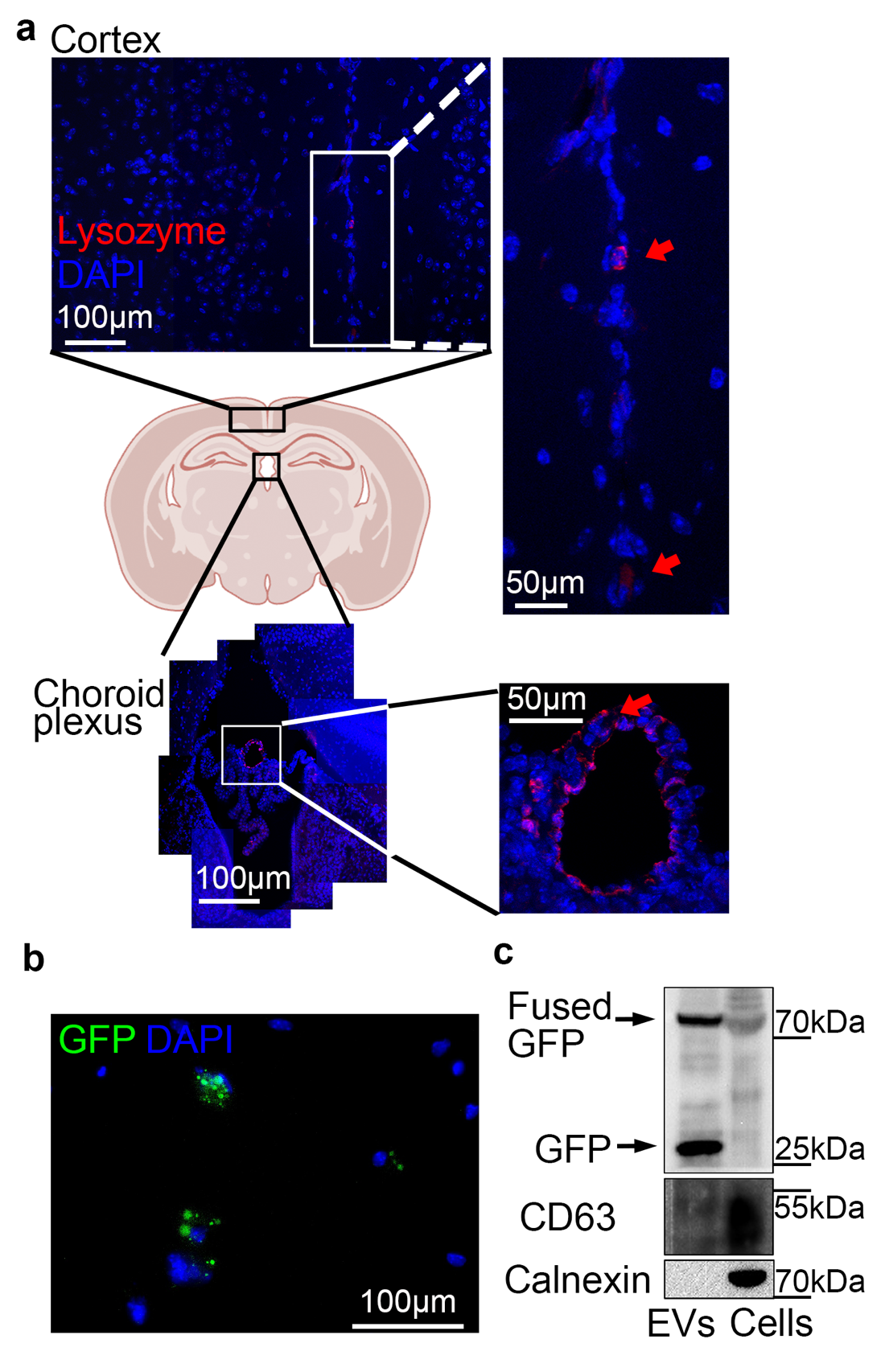

**Supplementary Fig. 2** Validation of the Lyz2^Cre^;CD63-emGFP^l/s/l^ reporter mouse for endogenous labeling of monocyte-derived extracellular vesicles.

a, Representative confocal images showing Lysozyme-positive myeloid cells (red) in the cerebral cortex and choroid plexus of Lyz2^Cre^;CD63-emGFP^l/s/l^ mice. Lysozyme-positive cells were readily detected within the meningeal/vascular interfaces and choroid plexus but were largely absent from the brain parenchyma, consistent with the restricted localization of peripheral monocyte-lineage cells under physiological conditions. Boxes indicate regions shown at higher magnification. Red arrows indicate representative Lysozyme-positive cells.

b, Representative fluorescence image of peritoneal monocytes of Lyz2^Cre^;CD63-eGFP^l/s/l^ mice demonstrating GFP-positive EV puncta signals.

c, Immunoblot characterization of EV preparations isolated from Lyz2^Cre^;CD63-eGFP^l/s/l^ monocytes. EVs expressed GFP and the canonical EV marker CD63 but lacked the endoplasmic reticulum marker calnexin, confirming successful labeling and isolation of monocyte-derived EVs.

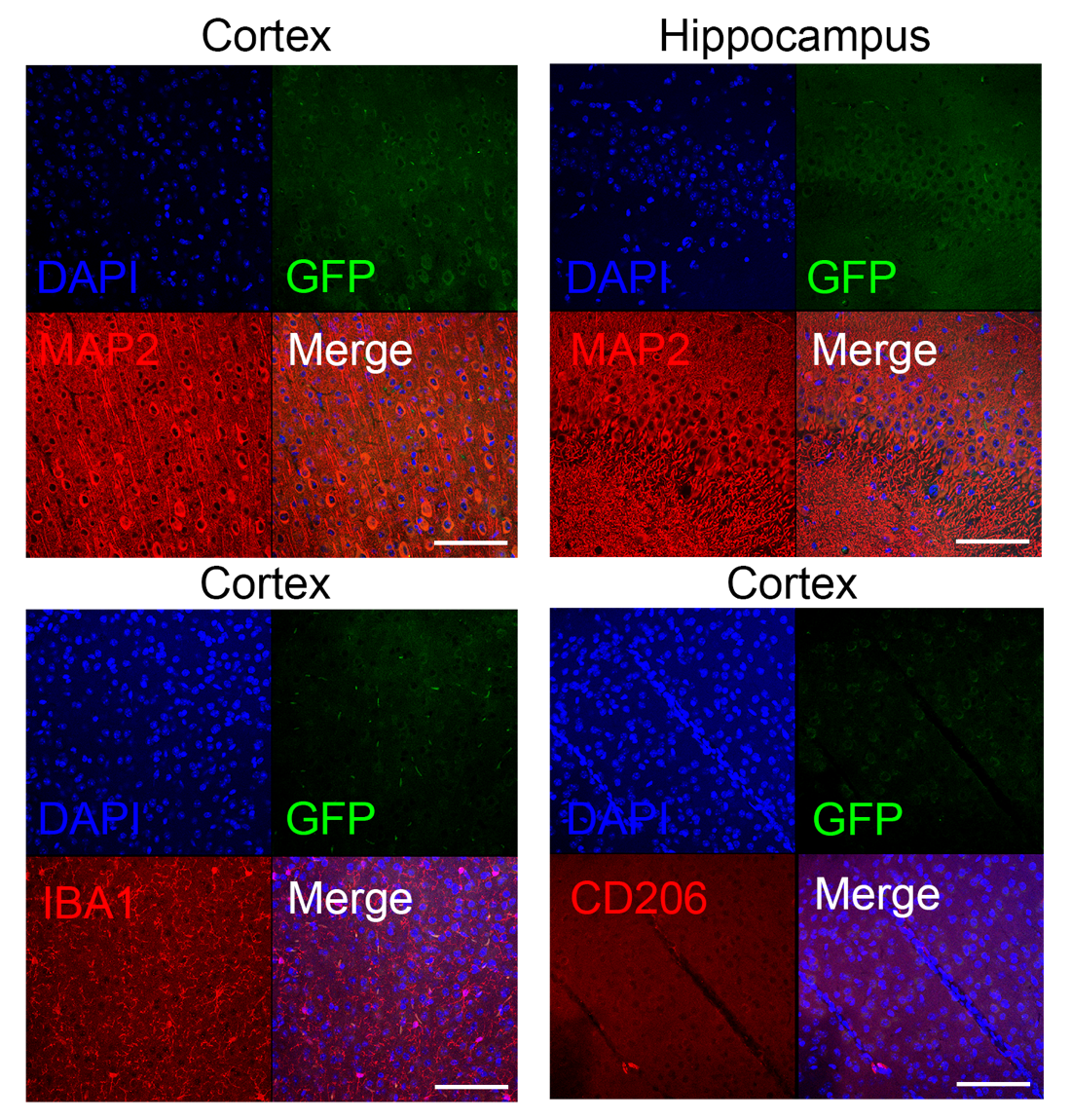

**Supplementary Fig. 3 Wild-type littermates exhibit no detectable CD63-GFP fluorescent signal in the brain.**

Representative confocal images of the somatosensory cortex and hippocampus from wild-type littermate mice processed under identical imaging and acquisition settings as the Lyz2^Cre^;CD63-emGFP^l/s/l^ reporter mice. Brain sections were stained for MAP2, IBA1, CD206 (red) and counterstained with DAPI (blue). No GFP fluorescent signal was detected in either brain region, confirming the specificity of GFP signals observed in Lyz2^Cre^;CD63-emGFP^l/s/l^ mice. Scale bars, 50 μm.

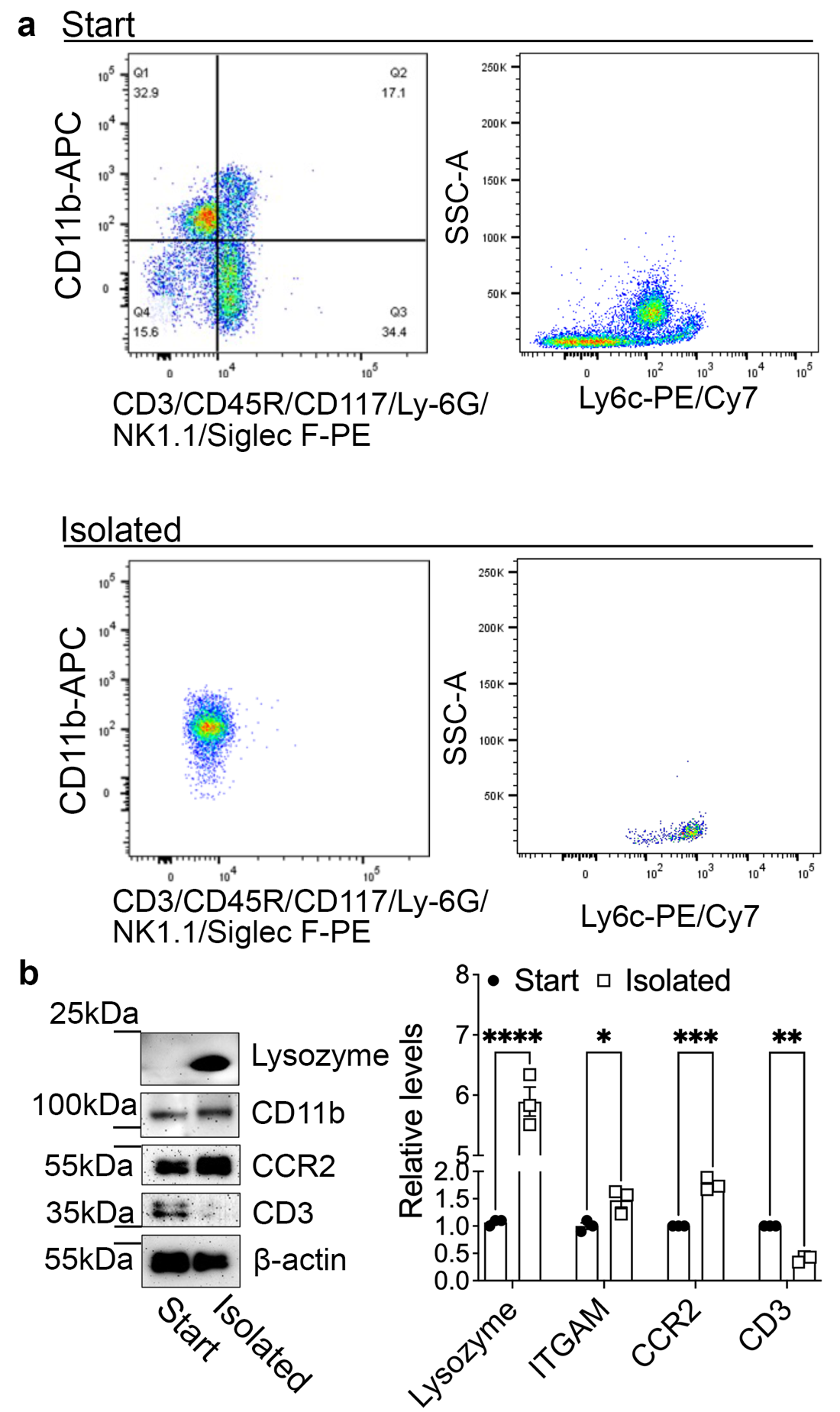

**Supplementary Fig. 4. Characterization of primary peripheral monocytes isolated from mouse whole blood.**

a, Representative flow cytometry analysis of whole blood before (Start) and after magnetic bead isolation (Isolated). Isolated cells were enriched for CD11b⁺Ly6C⁺ monocytes with minimal Ly6G⁺ neutrophil contamination.

b, Representative immunoblot analysis of whole blood (Start) and isolated peripheral monocytes (Isolated). Monocyte markers Lysozyme, ITGAM (CD11b), and CCR2 were enriched following isolation, whereas the T-cell marker CD3 was markedly reduced. β-actin served as a loading control.

Data are presented as mean ± s.e.m. Statistical analyses are described in the Methods.

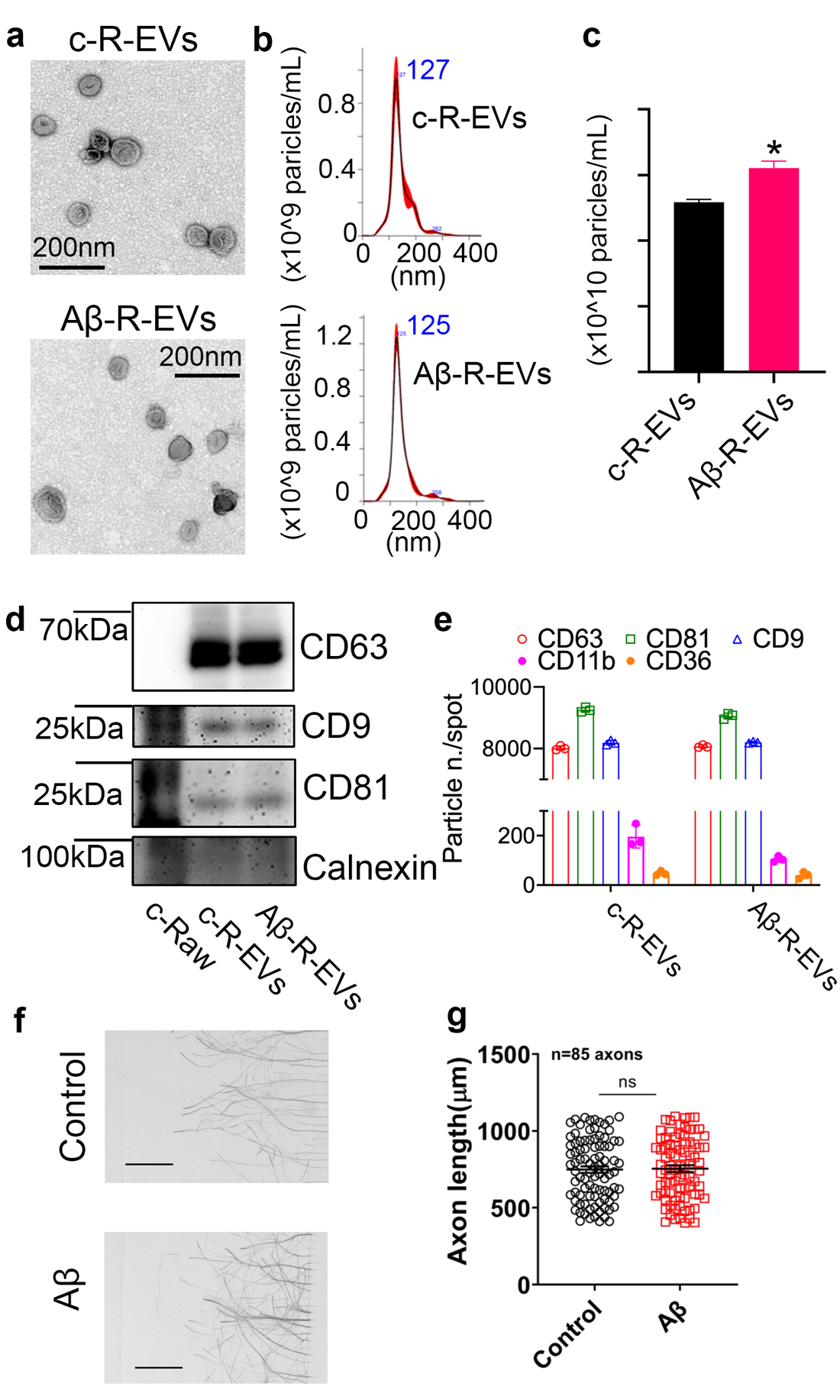

**Supplementary Figure 5 Characterization of EVs released from Aβ-stimulated monocytes.**

a, Representative TEM images of EVs isolated from control RAW264.7 monocytes (c-R-EVs) and Aβ₁₋₄₂-stimulated monocytes (Aβ-R-EVs).

b, NTA showing size distributions of c-R-EVs and Aβ-R-EVs. Peak particle diameters are indicated.

c, Quantification of EV particle concentration demonstrating a modest increase in EV release following Aβ stimulation.

d, Immunoblot characterization of EV preparations. EVs expressed the canonical EV markers CD63, CD9, and CD81, while lacking the endoplasmic reticulum marker calnexin, confirming successful EV isolation.

e, Single-particle phenotyping (ExoView) demonstrating comparable expression of canonical EV markers (CD63, CD81, and CD9) together with monocyte-associated surface markers (CD11b and CD36) on c-R-EVs and Aβ-R-EVs, indicating that Aβ stimulation increases EV production without substantially altering EV identity.

f, Representative images of primary cortical axons following direct treatment with 100 nM oligomeric Aβ_1-42_ or vehicle control.

g, Quantification of axonal length showing that direct exposure to 100 nM oligomeric Aβ_1-42_ did not significantly affect axonal outgrowth under these experimental conditions.

Scale bars in f, 200 μm. Data are presented as mean ± s.e.m. Statistical analyses are described in the Methods.

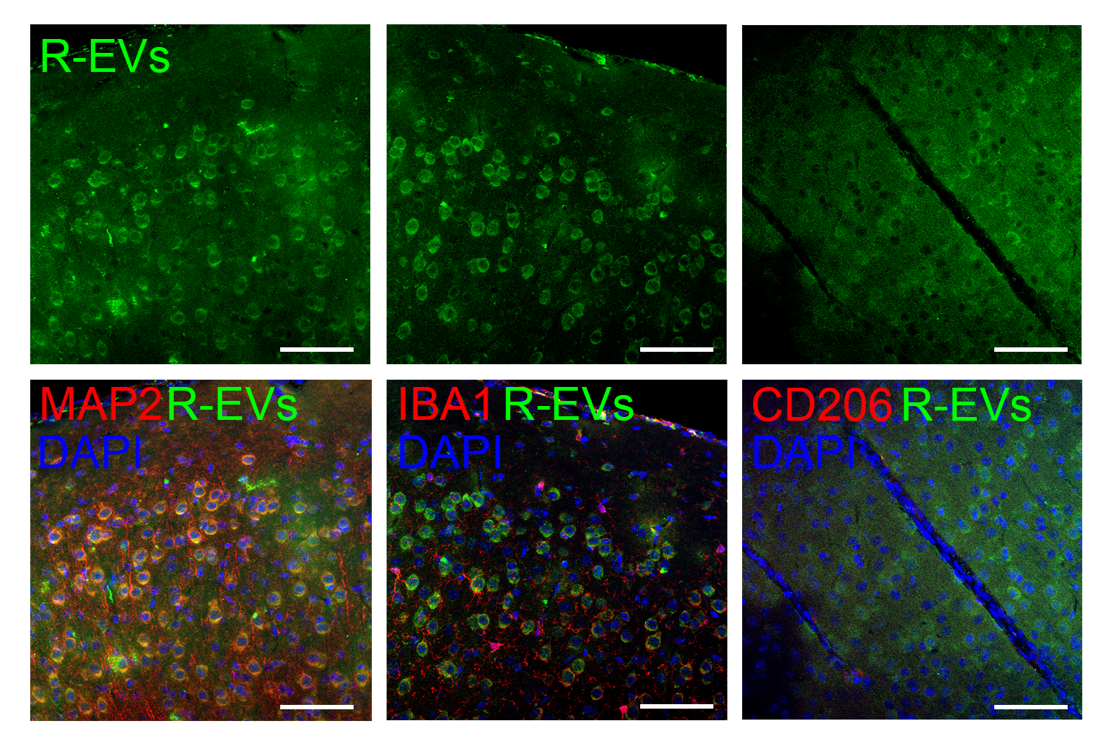

**Supplemental Fig. 6 Systemically administered monocyte-derived EVs are present in the mouse brain.**

Representative confocal images of brain sections collected following systemic administration of fluorescently labeled RAW264.7-derived EVs (R-EVs). Positive EVs were detected throughout the brain parenchyma and were frequently observed in close proximity to MAP2-positive neurons, whereas limited spatial association was observed with IBA1-positive microglia or CD206-positive perivascular macrophages. Scale bars, 50 μm.
